# Investigating the mechanisms underlying Bortezomib resistance

**DOI:** 10.1101/2023.07.29.551081

**Authors:** Kalliopi Zafeiropoulou, Georgios Kalampounias, Spyridon Alexis, Daniel Anastasopoulos, Argiris Symeonidis, Panagiotis Katsoris

## Abstract

Proteasome inhibitors such as Bortezomib, represent an established type of targeted treatment for several types of hematological malignancies, including multiple myeloma, Waldenstrom’s macroglobulinemia and mantle cell lymphoma, based on the cancer cell’s susceptibility upon impairment of the proteasome-ubiquitin system. However, a major problem limiting their efficacy is the emergence of resistance. Their application on solid tumors is currently being studied, while simultaneously, a wide spectrum of hematological cancers, such as Myelodysplastic Syndromes show minimal or no response to Bortezomib treatment. In this study, we utilize the prostate cancer cell line DU-145 to establish a model of Bortezomib resistance, studying the underlying mechanisms. Evaluating the resulting resistant cell line, we observed restoration of proteasome chymotrypsin-like activity, regardless of drug presence, an induction of pro-survival pathways, and the substitution of the Ubiquitin-Proteasome System role in proteostasis by induction of autophagy. Finally, an estimation of the oxidative condition of the cells, indicated that the resistant clones reduce the generation of reactive oxygen species induced by Bortezomib, to levels even lower than those induced in non-resistant cells. Our findings elucidate key proteins of survival and stress regulation pathways as potential pharmaceutical targets, which could increase the efficiency of the proteasome-targeting therapies, thus expanding the group of molecular targets for neoplastic disorders.

## Introduction

The Ubiquitin–proteasome pathway is the most important intracellular proteolytic system, and the integrity of its function is crucial for cell homeostasis (1). Proteasome substrates encompass signaling molecules, tumor suppressors, cell-cycle regulators, transcription factors, inhibitory molecules (whose degradation activates other proteins), and anti-apoptotic proteins (e.g., Bcl-2), among others (2). When degradation of these proteins is blocked, the detrimental effect is potentially enormous, especially for rapidly dividing cancer cells, which require increased availability of growth-promoting proteins, to sustain the accelerated and uncontrolled rate of mitosis, characteristic of cancer cell development and spread (3). Consequently, inhibition of the proteasome may delay cancer progression, by interfering with the regular degradation of cell-cycle proteins. Indeed, the inhibition of proteasomal function results in the induction of programmed cell death in several cell lines (apoptosis) (4–7), and therefore, proteasome inhibitors have been characterized as potential anticancer drugs.

Bortezomib, initially designated as PS-341 (Velcade®), reversibly inhibits the chymotrypsin-like activity of the proteasome. Chemically, Bortezomib is a dipeptidyl boronic acid analog derived from leucine and phenylalanine. It has been shown to inhibit tumor cell proliferation, adhesion, and metastasis in many in vivo and in vitro models, and has been approved by the US Food and Drug Administration (FDA) for use in cancer treatment (8). Several clinical trials have revealed that Bortezomib can be used to treat many types of solid tumors alone or in combination with other chemotherapeutic drugs. Bortezomib -first approved for patients with multiple myeloma-has also demonstrated inhibitory effects on colon-gastric cancer (9,10), breast cancer (11,12), prostate cancer (13,14), and lung cancer (15,16) as well as other cancer types.

Despite the impressive initial response, Bortezomib efficacy is limited by the rapid emergence of resistance. Almost 20 years after its approval, the mechanism of resistance to Bortezomib remains unclear. Several studies have been conducted to fully understand the mechanism underlying this resistance. Recently, some of the mechanisms of drug resistance have been elucidated. In vitro studies in lymphoma and leukemia cell lines have shown that Bortezomib resistance is developed either following mutations of the β5-subunit gene and/or β5 proteasome subunit gene overexpression (17,18) that leads to elevated chymotrypsin-like activity (19).

Normally, Bortezomib stabilizes p21, p27 and, p53, as well as some pro-apoptotic proteins (20,21) leading to tumor cell death. On the contrary, Bortezomib-resistant cells have been proven to evade apoptosis, by losing their ability to stabilize and accumulate the pro-apoptotic proteins (22). However, despite the importance of cell cycle regulation in cancer, there is little information about the precise role of cell cycle regulators in the development of Bortezomib resistance. Additionally, resistance has been associated with elevated phosphorylation levels of pro-survival proteins belonging to MAPKs, PIP3/AKT/mTOR and JAK/STAT pathways, and the underlying crosstalk (11,23,24). Key molecules of these pathways have been reported to be activated inside the resistant cells, elucidating the importance of survival protein modulation (25). Furthermore, upregulation of pathways that suppress apoptosis and induce autophagy (26,27) has also been reported in bortezomib-resistant cells. It is well known that these two mechanisms’ interplay is very crucial for cell survival. Autophagy blocks the activation of apoptosis and in turn, apoptosis blocks the activity of autophagy, through caspase-mediated cleavage of the autophagic proteins (28). Autophagy refers to pathways by which abnormal, malfunctioning, or simply excessive cytoplasmic material is delivered into the lysosomes for degradation (29). Autophagy is linked to the UPS (30), regulating proteostasis, and partially substituting it (31,32). Many multidrug-resistant types of cancers have been reported to possess upregulated autophagy biomarkers (33). In parallel, p62 and LC3 are elevated and an important role has been assigned to Beclin-1, which regulates the autophagy-apoptosis equilibrium (34).

Some latest reports indicate ROS-induced cell cycle arrest as a mechanism of drug resistance (35). However, until today, there is no clear evidence if ROS levels are crucial for the establishment of Bortezomib resistance although modulating intracellular ROS levels appears to be crucial for overcoming multidrug resistance in cancer cells (36). ROS levels may affect the phosphorylation of these cell cycle regulators and hence influence cell cycle progression. Thus, G1-arrested melanoma cells, were resistant to apoptosis, induced by the proteasome inhibitor bortezomib, irrespective of the factor mediating the arrest, a finding suggesting that induction of G1 arrest may result in Bortezomib resistance (37).

In the present study, we established a bortezomib-resistant prostate cancer cell line, DU-145 RB60, to study the differential effects of the drug in naïve (DU-145) and bortezomib-resistant (DU-145 RB60) cells. Hereby, we focused on changes in the accumulation of β5 subunits (PSBM5), poly-ubiquitinated proteins, cell cycle and autophagy regulators, as well as on the apoptotic rate and ROS levels in both cell populations.

## Materials and Methods

### Reagents and materials

Cell culture medium RPMI 1640 and all other culture reagents were purchased from Biowest (France). The specific proteasome inhibitor, Bortezomib (formerly known as Velcade^TM^), was generously provided by Janssen-Cilag (Greece). All culture plates were purchased from Greiner Bio-One (Austria).

### Cell Culture

The human prostate cancer epithelial cell line DU-145 (ATCC) was cultured in RPMI 1640 medium, supplemented with 10% Fetal Bovine Serum (FBS), 100 units/ml penicillin, and 100 μg/ml streptomycin. Cultures were incubated at 5% CO_2_ and 100% humidity at 37°C.

### Cell viability assay

To assess viability, cells were plated at a density of 50.000 cells per well inside 24-well plates. After cell spreading and adhesion, they were treated with the indicated concentrations of Bortezomib in RPMI 1640 supplemented with 10% FBS for 24 to 72 h. The inhibitory effect of Bortezomib on cell growth was measured using the crystal violet assay (38). Adherent cells were fixed with methanol and stained with a 0.5% crystal violet in 25% methanol aqueous solution for 20 min. After gentle rinsing with water, the retained dye was extracted using a 30% acetic acid aqueous solution, and the absorbance was measured at 595 nm using a plate reader spectrophotometer.

### Microchemotaxis/Transwell Chambers

Known numbers of cells were transferred inside Transwell chambers/inserts containing serum-free medium with or without Bortezomib. The inserts were then submerged inside wells of 24-well plates containing medium supplemented with 20% FBS with or without Bortezomib. The cells were left to migrate for an indicated interval and then were washed with PBS solution and fixed with methanal solution. The fixed cells were stained with 0.33% toluidine blue solution and photographs were taken on a photonic microscope at an x20 magnification (38).

### Wound Healing Assay

Cells were grown on 6-well plates until the formation of a confluent monolayer. Subsequently, cells were scratched in a cross-like manner using a tip and the medium was replaced with Bortezomib-containing RPMI 1640 supplemented with 10% FBS. Photographs were taken immediately after the scratches/wound formation as well as after key time points (39).

### Proteasome activity assay

Cells were exposed to a range of various Bortezomib concentrations for 24 h. Total proteins were extracted from the cells, using sonication and incubation in a solution containing 50 mM ΗΕPES, 20 mM KCl, 5 mM MgCl_2·_H_2_O, 1 mM Dithiothreitol (DTT) (Cat. 10197777001, Merck) with a pH value of 7.81. The extract was incubated with proteasome fluorogenic substrate-peptide LLVY-AMC (Suc-Leu-Leu-Val-Tyr-7-amide-4-methylcoumarin) (Cat. Number 3120-v, Peptide Institute Inc.), with or without the proteasome inhibitor MG-132 (Cat. Number 3175-v, Peptide Institute Inc.) inside flat-bottom black polystyrene 96-well plate for fluorescence for 1 h at 37°C (40). Fluorescence intensity was measured at 380 nm excitation wavelength and 460 nm emission wavelength. Total protein concentration was estimated using the Q5000 Nanodrop Quawell spectrophotometer.

### Western blot analysis

Cells were starved from any drugs/inhibitors for 24 h, and then incubated with the indicated concentrations of Bortezomib for varying times. Subsequently, they were washed twice with PBS 6 solution and lysed using RIPA buffer. Total proteins were determined using the Bradford Assay. Equal amounts of total proteins were mixed with Laemmli’s Sample Buffer 2X solution containing 5% β-ME, and the samples were denatured at 95°C for 10 min. Proteins were separated by SDS-PAGE and transferred to an Immobilon-P membrane (Millipore, USA) for 30 min using the Towbin’s transfer buffer in a semi-dry transfer system. The membrane was blocked in TBS containing 5% skimmed milk and 0.1% Tween-20 for 1 h at 37°C. Membranes were then probed with primary antibodies (Table 1) overnight at 4°C, under continuous agitation.

**Table 1.** List of primary antibodies used for Western blot analyses.

| Antibody | Supplier | Dilution |
| --- | --- | --- |
| **b-actin antibody (monoclonal mouse)** | CST #3700 | 1: 2000 |
| **Proteasome 20S β5 subunit antibody (polyclonal mouse)** | Santa Cruz #sc-393931 | 1: 1000 |
| **Ubiquitin antibody (P4D1) (monoclonal mouse)** | Santa Cruz #sc-8017 | 1:2000 |
| **p21 antibody (monoclonal rabbit)** | CST #2947 | 1:1000 |
| **p27 antibody (monoclonal rabbit)** | CST #3686 | 1:1000 |
| **p53 antibody (monoclonal mouse)** | Invitrogen # MA5-12557<br>(ThermoFisher Scientific) | 1:500 |
| **p44/42 MAPK (Erk1/2) Antibody (polyclonal rabbit)** | CST #9102 | 1:1000 |
| **Phospho-p44/42 MAPK (Erk1/2) (Thr202/Tyr204) Antibody (polyclonal rabbit)** | CST #9101 | 1:1000 |
| **Phospho-p38 MAPK (Thr180/Tyr182) antibody (monoclonal rabbit)** | CST #9215 | 1:1000 |

The blot was then incubated with the appropriate secondary antibodies (Anti-rabbit IgG Antibody, CST#7074, or Anti-mouse IgG antibody, CST# 7076) (both diluted 1:2000) coupled to horseradish peroxidase, and bands were detected with the SuperSignal™ West Femto Maximum Sensitivity Substrate (Thermo Scientific™ #34096), according to the manufacturer’s instructions. Where indicated, blots were stripped in buffer containing 62.5 mM Tris HCl pH 6.8, 2% SDS, 100 mM 2-mercaptoethanol for 30 min at 50°C and reprobed with primary antibodies. Quantitative estimation of band size and intensity was performed through analysis of digital images using ImageJ (41).

### Immunofluorescence confocal microscopy

Cells were cultured on glass coverslips (MGF-slides, Germany) for 24 h and then incubated with appropriate Bortezomib concentrations. After this interval, they were gently rinsed with PBS and fixed in 4% paraformaldehyde aqueous solution for 15 min at room temperature. Subsequently, they were rinsed three times with PBS, permeabilized for 15 min in PBS containing 0,1% Triton X-100 and blocked in PBS containing 5% skimmed milk for 1 h at room temperature. Cells were then incubated for 1 h, at room temperature, with the primary antibodies/fluorophores (Table 2).

**Table 2.** List of primary antibodies used for immunocytochemical staining.

| Antibody | Dilution |
| --- | --- |
| **beta-actin antibody (monoclonal mouse)** | 1: 1000 |
| **p21 antibody (monoclonal rabbit)** | 1:500 |
| **p27 antibody (monoclonal rabbit)** | 1:500 |
| **DAPI** | 0,1 µg/ml |

After rinsing with PBS Tween-20, the coverslips were incubated with secondary antibodies; Donkey anti-Rabbit IgG (H+L) Highly Cross-Adsorbed antibody, Alexa Fluor™ 647 (Invitrogen, #A-31573) or Goat anti-Mouse IgG (H+L) Cross-Adsorbed secondary antibody, Alexa Fluor™ 488 (Invitrogen, #A-11001) (Dilution 1:500) in permeabilization buffer. After rinsing three times in PBS, cells were mounted using Mowiol 4-88® (Sigma). To detect autophagy, live cells grown on coverslips were incubated with Lysotracker RED (Invitrogen) for 15 min and then immediately imaged. Labelling was performed using Leica Confocal Imaging System confocal and photographs were taken using the LAS X software. Lysotracker staining of live cells was also used to quantify the acidic protein content using Flow Cytometry.

### Flow Cytometry

Cells were incubated with the indicated concentrations of Bortezomib in RPMI 1640, supplemented with 10% FBS for appropriate time intervals (24-48 h). Subsequently, they were stained with Annexin-V conjugated with FITC (Invitrogen™ #A13199) for 20 min at room temperature to detect apoptosis (42). Propidium Iodide (PI) (BD Pharmingen^TM^ #556463) was used both as a necrotic marker during Annexin V assay, as well as during the cell cycle analysis to quantify DNA content. Staining with Lysotracker RED was optimized for acidic protein content estimation. The cells were stained for 45 min at 37° C using the concentration suggested for immunofluorescence and subsequently rinsed and analyzed. Finally, the cell-permeant 2’,7’-dichlorodihydrofluorescein diacetate (H_2_DCFDA) (Invitrogen, # D399), was used to estimate Reactive Oxygen Species (ROS) in these cells. The samples were analyzed in a FACS Calibur cytometer (BD, Biosciences). For each sample, 200,000 ungated events were acquired.

## Results

### Generation of a DU-145 Bortezomib-resistant clone

Bortezomib-resistant cells were established, through a gradual increase in the concentration of Bortezomib for at least 24 weeks, from 5 to 60 nM Bortezomib, and then maintained in the same concentration constantly for 12 weeks. The resistant clone was named DU-145 RB60 and it was cultured in parallel with the naïve DU-145 clone. The two clones were examined for Bortezomib-induced growth inhibition. Dose-response viability/proliferation curves showed 5-fold higher resistance (IC_50_: 60 nM) in DU-145 RB60 compared to the naïve DU-145 after 72 h of treatment with Bortezomib (Fig 1)**Error! Reference source not found.**. In addition, the DU-145 RB60 clone was evaluated for potential cross-resistance to a newer proteasome inhibitor, Carfilzomib, but this was of minor significance (Table 3).

**Fig 1.**
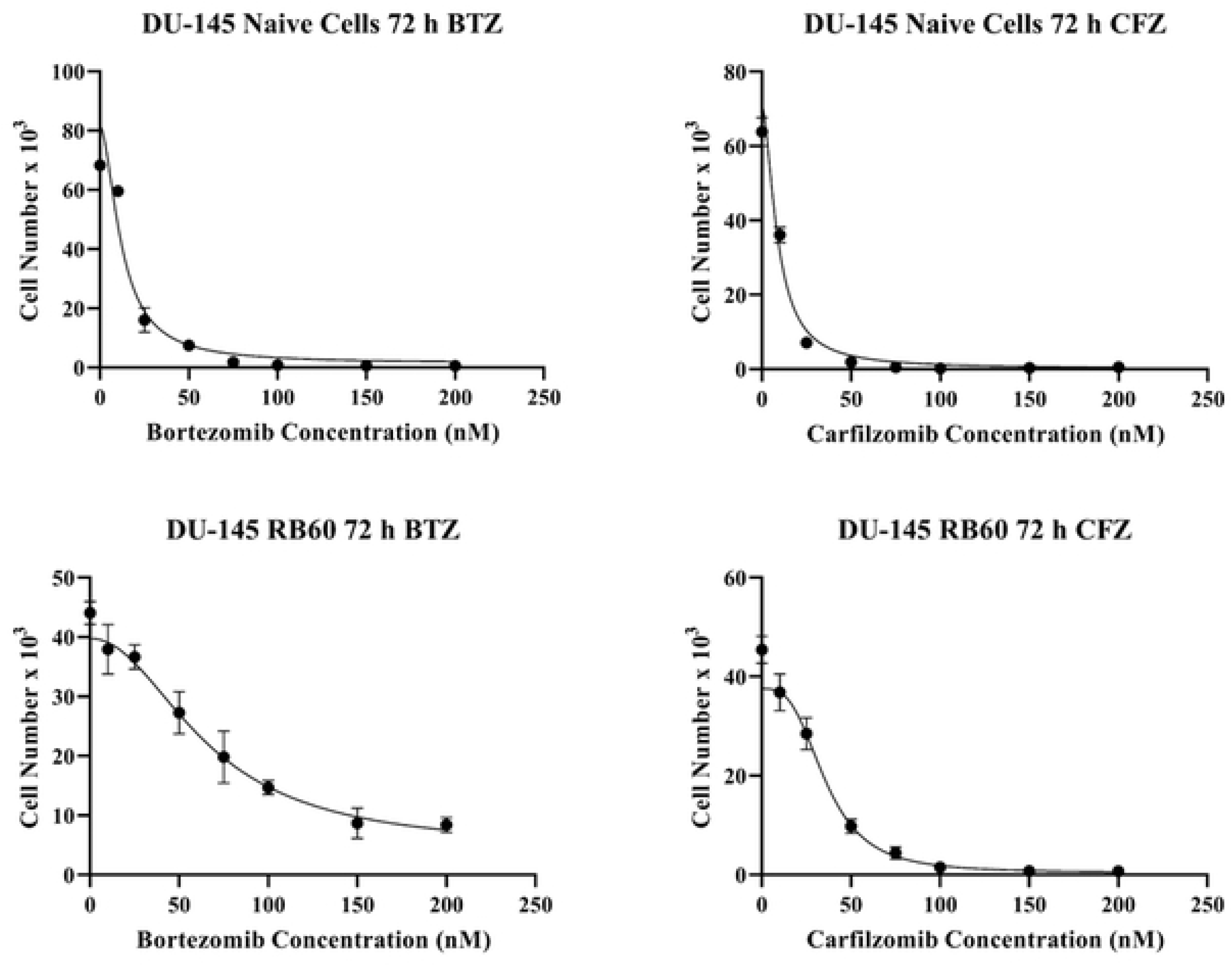
Proliferation curves of DU-145 naive and DU-145 RB60 cells. Equal numbers of cells were seeded on 24-well plates and after 24 h of attachment, proteasome inhibitors were added. Following 72 h of incubation, the cells were fixed and stained with crystal violet. The proliferation rate of naïve DU-145 and DU-145 RB60 resistant cells is assessed under the presence of Bortezomib **(A, C)** and Carfilzomib, respectively **(B, D)** indicating a 5-fold increase of the resistant cells’ ability to survive, compared to the naïve clone. A partial cross-resistance to Carfilzomib is also observed.

**Table 3.** IC_50_ calculation of naïve DU-145 and DU-145 RB60 cells. The data from crystal violet assays were analyzed using GraphPad Prism 8 and the calculated IC50 values are presented here. The resistant cells exhibited a 5-fold increase of Bortezomib tolerance while at some extent, cross-resistance was observed; the DU-145 RB60 clone achieved 4-fold Carfilzomib resistance compared to the naïve clone.

|  | IC<sub>50</sub> (nM) |  | IC<sub>50</sub> ratio |
| --- | --- | --- | --- |
| Cell type | DU-145 Naïve | DU-145 RB60 |  |
| Proteasome inhibitor |  |  |  |
| Bortezomib | 12.19±1.06 | 65.87±1.96 | 5.40 |
| Carfilzomib | 8.84±1.17 | 35.74±1.56 | 4.04 |

The baseline biological activities such as migration, chemotactic movement, and wound healing of the resistant clone DU-145 RB60 were also assessed (Figs 2 and 3). To study the effect of Bortezomib on migration, we incubated DU-145 RB60 cells with or without the presence of the drug in micro-chemotaxis chambers. The presence of 60 nM Bortezomib in the upper compartment did not affect the DU-145 RB60 clone’s ability to migrate. When we added 60 nM Bortezomib to the chemoattractant medium (lower compartment) naïve DU-145 cells failed to get mobilized as efficiently as the resistant clone, suggesting that the drug might act as a chemorepellent only for the naïve cells. Further dose escalation (to both migration and chemotaxis assays) revealed that doses up to 240 nM were required to achieve the same effects on the resistant clone. Finally, we assessed the wound-healing ability of both clones, with or without the presence of Bortezomib and we found that only the resistant clone achieved healing of the wound without changes, as compared to the control. Thus, resistant DU-145 RB60 cancer cells appeared to maintain all the main biological characteristics in the presence of Bortezomib, without significant changes compared to the naïve untreated DU-145 clones.

**Fig 2.**
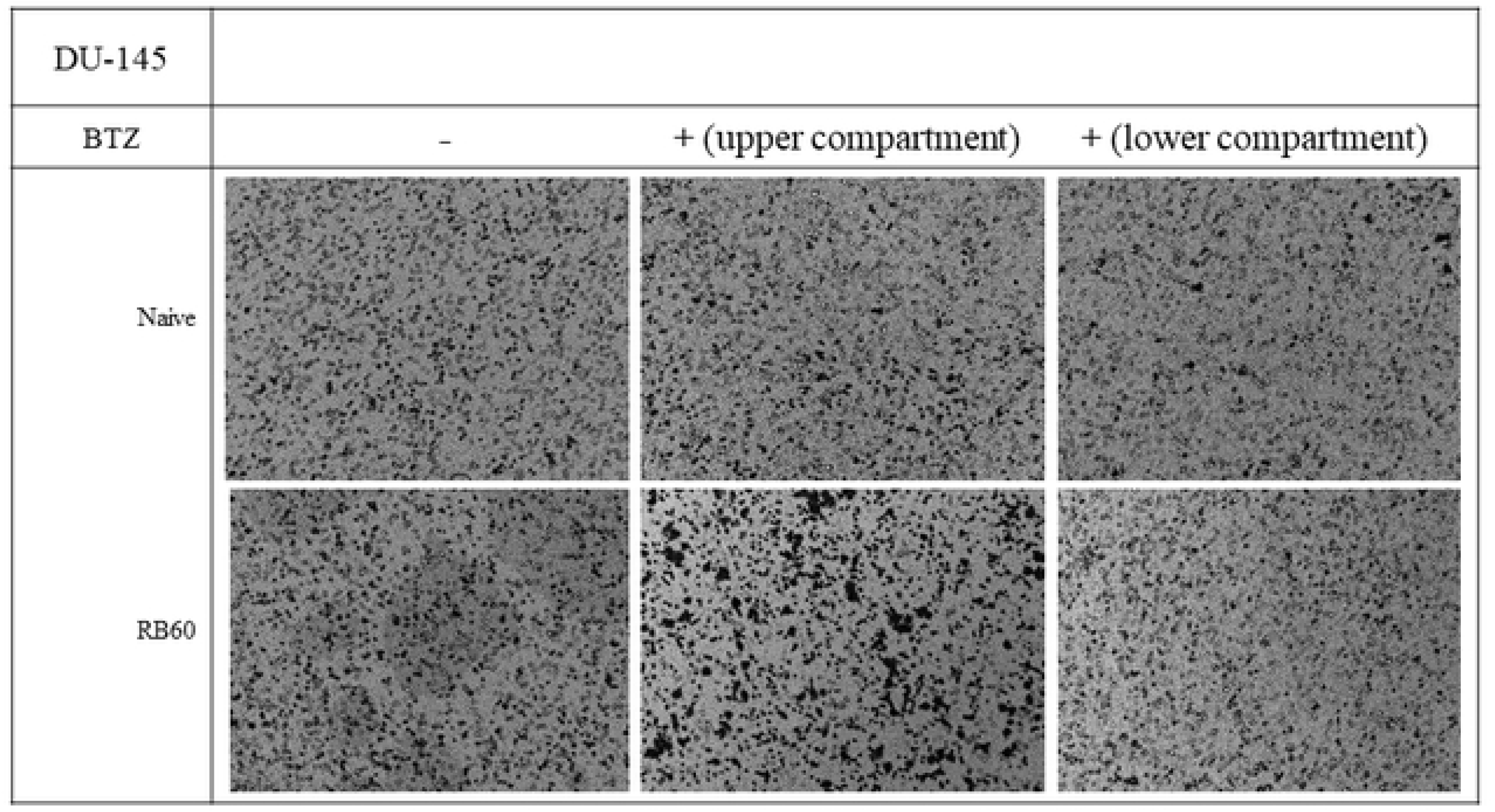
Chemotactic assay of DU-145 naive and DU-145 RB60 cells using Boyden chambers. Cells were transferred inside a chamber containing serum-free medium with or without Bortezomib. The chambers were placed inside microplates’ wells containing medium supplemented with 20% FBS and left to migrate for 24 h. The DU-145 cells when exposed to Bortezomib (20 nM) decreased their migration rate. Inhibition of migration was also observed when Bortezomib was added to the lower compartment indicating a chemorepellent role. The DU-145 RB60 cells were also repelled by Bortezomib (60 nM) while the presence of the drug in the upper compartment induced migration towards the other side of the membrane where Bortezomib was absent.

**Fig 3.**
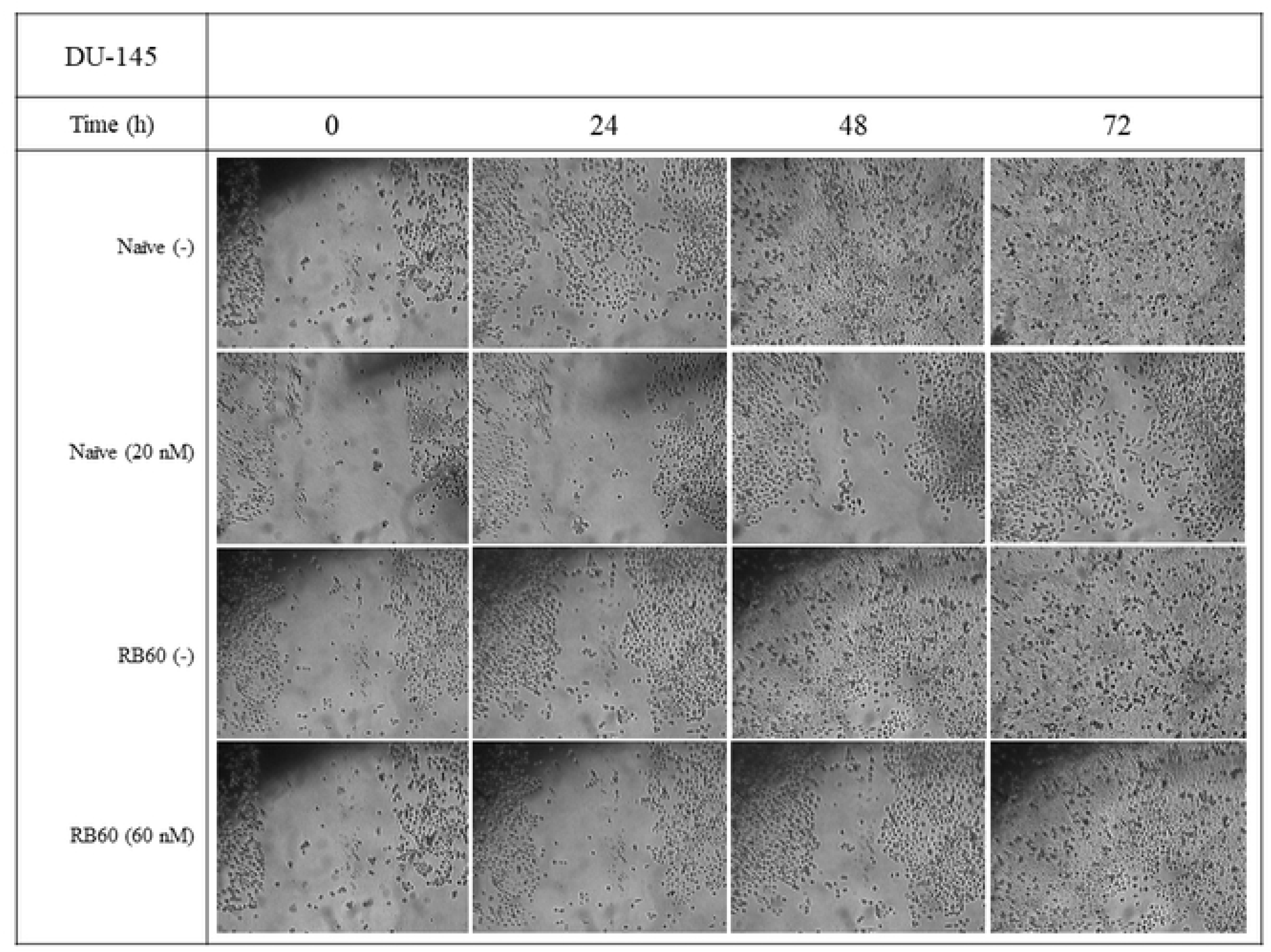
Wound healing assay of DU-145 naive and DU-145 RB60 cells. Cells were seeded on 6-well plates and left to form monolayers. After reaching the desired confluency, wounds were scratched, and the Bortezomib-free media were replaced with medium containing 10% FBS and Bortezomib. The naïve cells were assessed using 20 nM of Bortezomib and the DU-145 RB60 cells were assessed under the influence of 60 nM Bortezomib. Compared to the control group, the DU-145 naïve cells’ ability to heal wounds was heavily impaired by Bortezomib while the same effect was not observed on the DU-145 RB60 cells. The resistant cells were able to completely heal the scratches after 72 h of incubation and the same was achieved by the untreated naïve cells.

### Bortezomib-resistant clones successfully restore the UPS system activity, even on high drug doses

In cancer cells, Bortezomib successfully inhibits protein degradation and promotes the accumulation of polyubiquitinated proteins. To investigate the response of the resistant clone to Bortezomib, we assessed the polyubiquitination of the total proteins on both, resistant and naïve DU-145 clones by western blot analysis (Fig 4A). Time-course experiments showed that the naïve DU-145 clone accumulated polyubiquitinated proteins after incubation with 20 nM Bortezomib for 24 h, while on the DU-145 RB60 clone, the same effect was not observed (Fig 4B). The DU-145 RB60 cells accumulated ubiquitinated proteins only after incubation with >200 nM Bortezomib.

**Fig 4.**
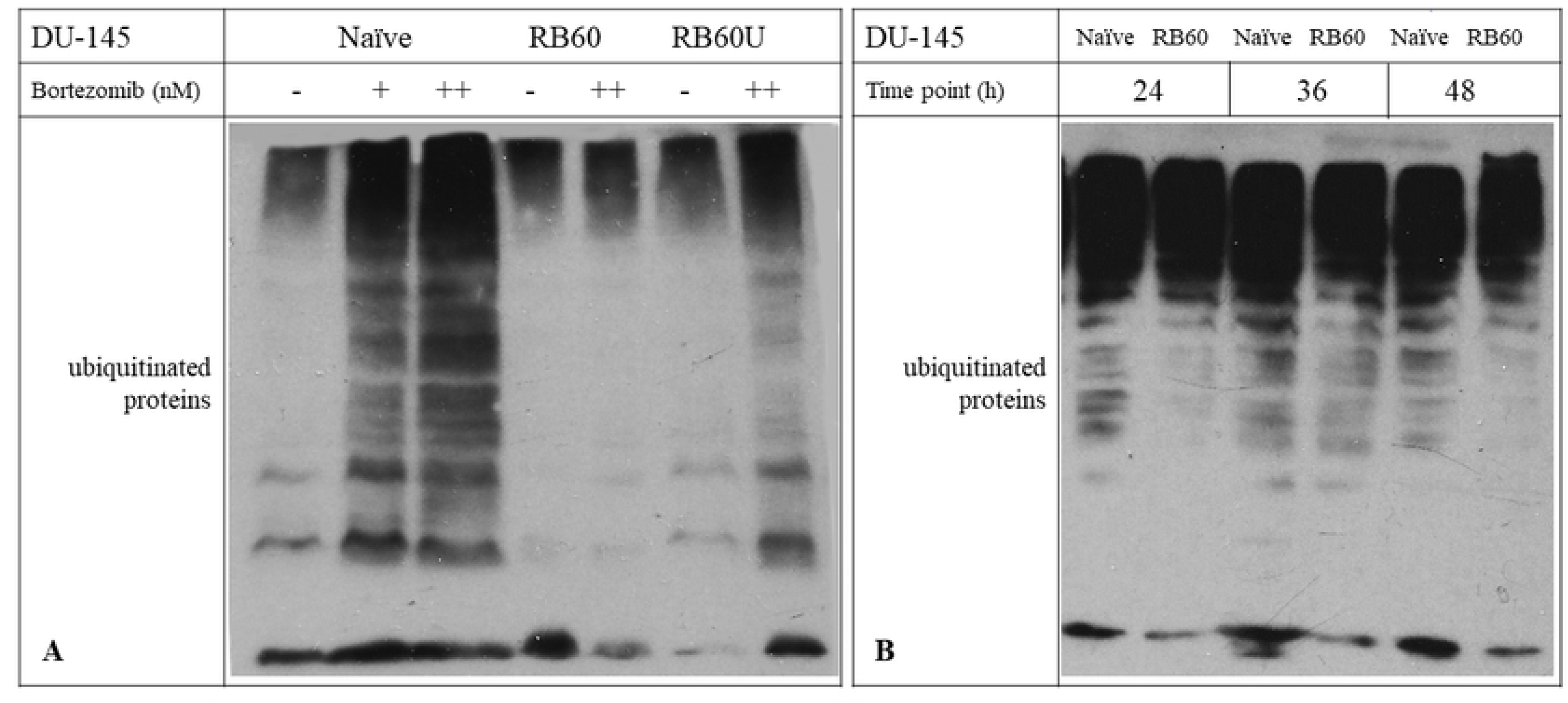
Western Blot analyses of ubiquitinated proteins in naive and resistant clones. **(A)** Dose-response experiments on Naive cells indicated accumulation of ubiquitinated proteins after 20 nM Bortezomib. The resistant cells studied did not accumulate ubiquitinated proteins following incubation with 60 nM of Bortezomib. The RB60U clone exhibited mild UPS impairment following a 60 nM Bortezomib treatment for 24 h. **(B)** Time-course experiments verified the stable ubiquitination levels at the key intervals of 24, 36 and 48 h following incubation with 20nM Bortezomib, validating the 24 h time point as an adequate moment to study effects on main signaling pathways.

This suggests that the resistant cells, despite their tolerance to the drug, exhibit an accumulation of ubiquitinated proteins at higher concentrations (S1 Fig.). Subsequently, we chose the 24 h time point and the dose of 60 nM Bortezomib to conduct the signaling investigation experiments. At this specific time point, the naïve cells were affected by the drug, whereas the resistant clone remained intact and the RB60U cells showed mild accumulation of polyubiquitinated proteins.

In many similar studies, the molecular mechanism of Bortezomib resistance relies on a dramatic overexpression of the PSMB5 protein (17). To clarify whether this mechanism might also be active in our resistant cancer cell line, we determined the expression of proteasome subunit β5 on DU-145 RB60 cells and the naïve cells by western blot analysis (Fig 5A). Overall, higher expression of this proteasome subunit was seen on the DU-145 RB60 cells, compared to the naïve DU-145 cells. Protein levels of the β5 subunit were higher (P < 0.01) on the DU-145 RB60 clone with or without Bortezomib. To clarify if this increased β5 subunit expression might lead to a subsequent induction of elevated proteasome activity we determined the basal chymotrypsin-like (ChT-L) activity on both, resistant and naïve clones. Our results showed that the ChT-L activity of the DU-145 RB60 clone (even after Bortezomib short-term deprivation) was significantly higher than that of the naïve DU-145 clone, suggesting the emergence of an acquired Bortezomib resistance, which is independent of the constant drug presence (Fig 5B).

**Fig 5.**
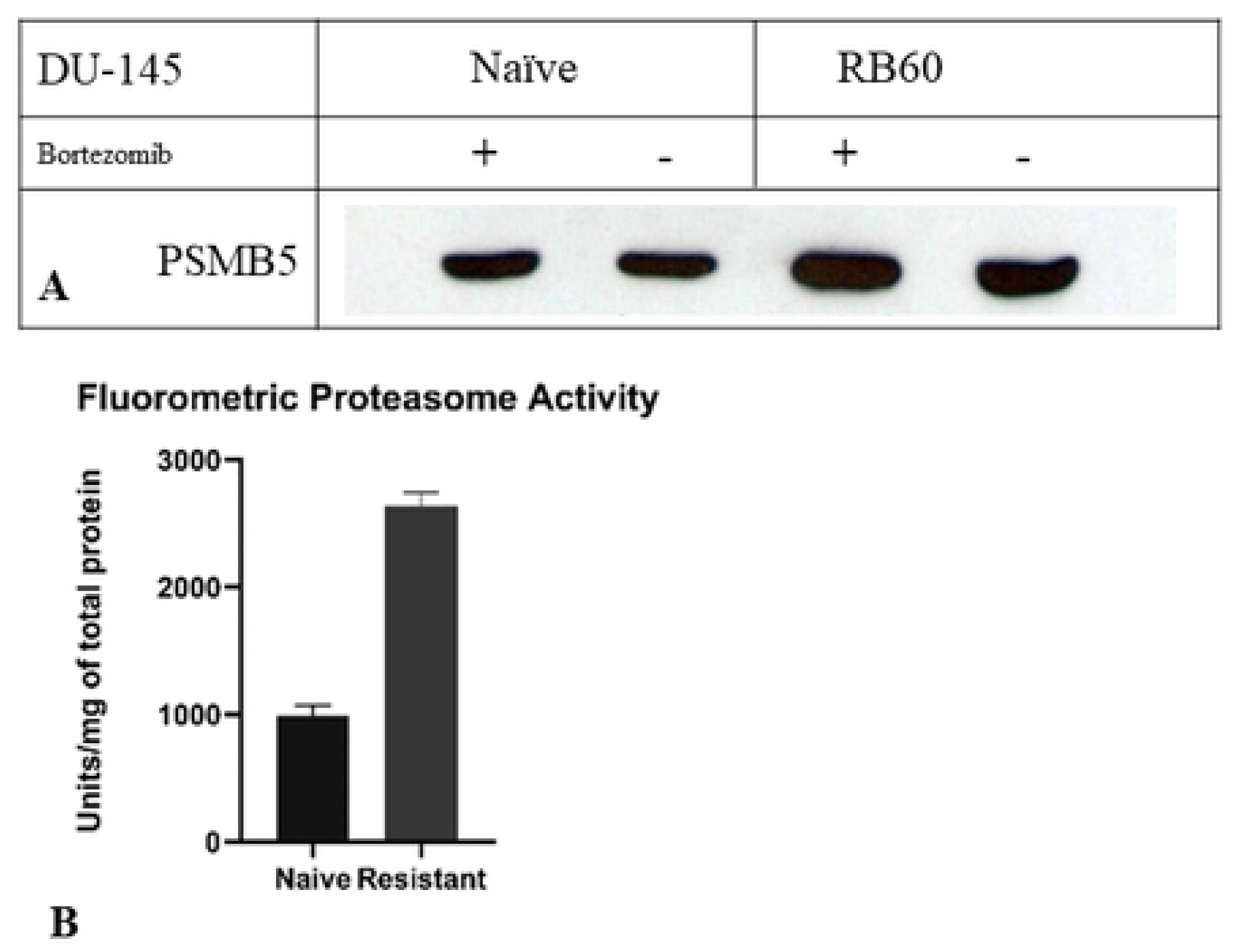
Proteasome activity assay; Chymotrypsin-like activity and PSMB5 accumulation. **(A)** PSMB5 (or β5) is the most ubiquitous proteasome subunit exhibiting ChT-L activity. Treatment with Bortezomib slightly increased PSMB5 expression on DU-145 cells while the resistant clone DU-145 RB60 exhibited permanent overexpression, inducible upon further treatment with Bortezomib. **(B)** The Chymotrypsin-like activity of DU-145 naïve and DU-145 RB60 cells was also assessed using fluorometry, showing a notable change in the resistant cells.

It was also crucial to clarify if this acquired phenotype is inherent or can be reversed after long-term drug withdrawal, so we maintained a subclone of the DU-145 RB60 deprived of Bortezomib for six months, to determine the stability of this characteristic and assess the effects of Bortezomib on the cells’ proliferation rate and changes on main signaling pathways. These long-term untreated cells were named DU-145 RB60U cell clone.

### Resistant cells evade apoptosis induced even by high Bortezomib doses

To investigate the mechanism by which the resistant clone achieves to avoid drug toxicity and survives in higher Bortezomib concentrations, we determined the apoptotic rate in both clones by flow cytometry using the Annexin V/Propidium Iodide assay. Incubation of naïve DU-145 cells with 60 nM Bortezomib led to the induction of apoptosis at a significant degree (P < 0.01) (Fig 6B), an effect that was not observed on the DU-145 RB60 clone (Fig 6D). The DU-145 RB60 clone appeared to be unaffected by low and medium doses of Bortezomib and exhibited the same apoptotic rate, as untreated naïve DU-145 cells (Figs 6A and 6C). Both, early and late apoptotic rates of DU-145 RB60 cells remained at least 6-fold lower than the respective of naïve cells, suggesting that these cells were capable to suppress cell death induction mechanisms and eventually evade apoptosis.

**Fig 6.**
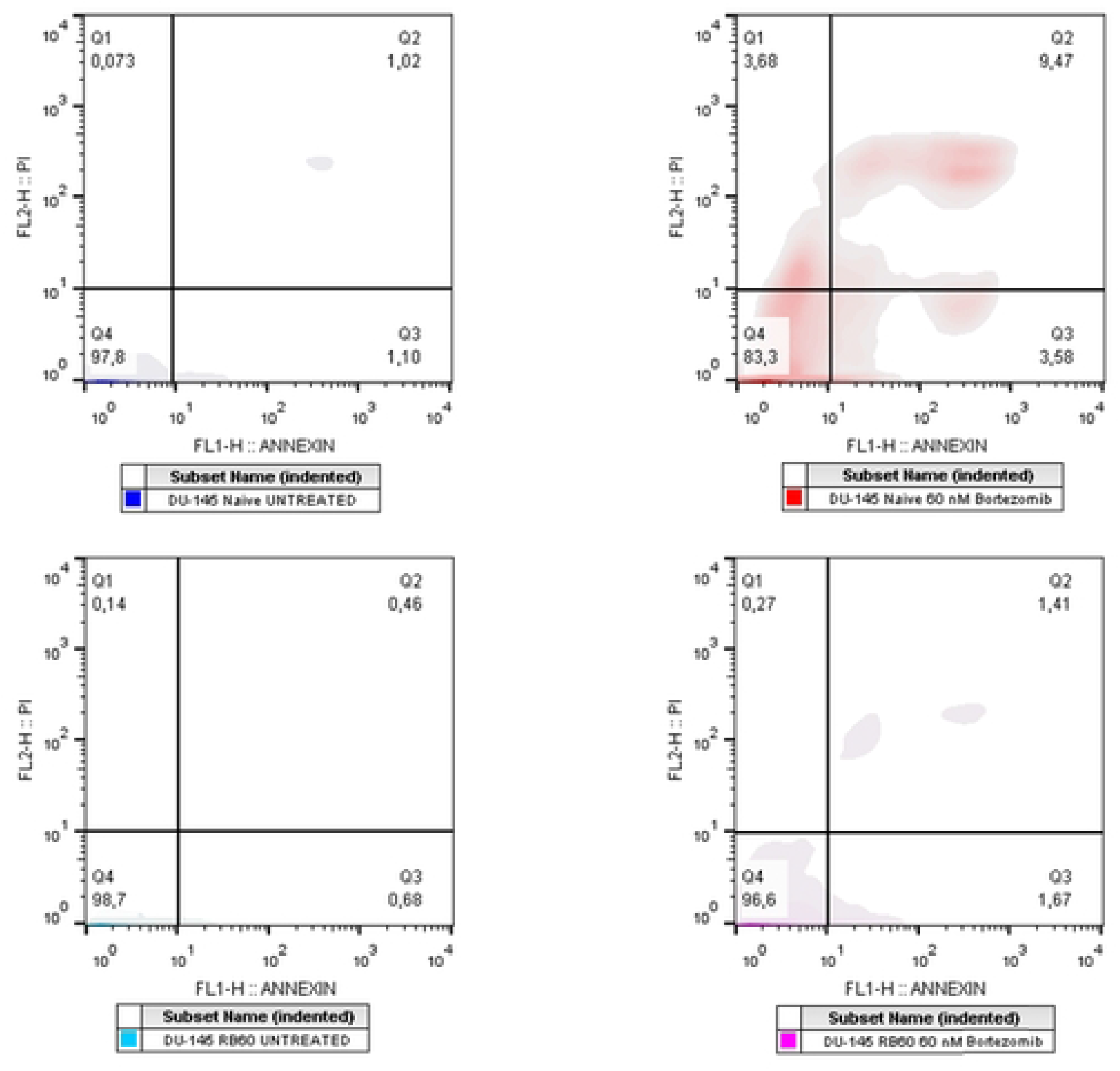
Apoptosis assay using Annexin V and PI. Flow cytometry analysis using Annexin V-FITC and PI indicated that the DU-145 RB60 cells (C) have fully restored their apoptotic rate to basal levels **(A)**. Cells were treated with Bortezomib for 48 h and subsequently trypsinized and stained with Annexin V/PI. Treatment with 60 nM of Bortezomib greatly induced apoptosis diminishing viability by 15% in the first 48 h on naïve DU-145 cells **(B)**, while the same treatment, only slightly increased the percentage of apoptotic DU-145 RB60 cells **(D)**. The evasion of apoptosis on DU-145 cells is only lost upon treatment with Bortezomib doses exceeding 180 nM.

### Resistant cells overthrow cell cycle arrest induced by Bortezomib

The effects of Bortezomib on the main apoptotic signaling molecules have been long-established (4–7). However, the consequences of the drug on cell cycle regulation and the interrelationship between those two mechanisms are not fully understood. In general, Bortezomib inhibits cell cycle progression mainly by dysregulating the turnover of various cell cycle regulators, such as cyclins and CDK-associated molecules (43,44).

The accumulation of p21^waf1/kip1^, a known cell cycle inhibitor, has been shown to correlate with Bortezomib’s action (6,20,45,46). Incubating the cells with Bortezomib led to an increase in the nuclear levels of p21 protein in a dose-response manner (Fig 7B) and in a time-course manner, which showed a peak after 24-36 h of incubation. It is well-known that The nuclear localization of p21 mainly causes cell cycle arrest, while its cytoplasmic presence has been associated with antiapoptotic activity (47).

**Fig 7.**
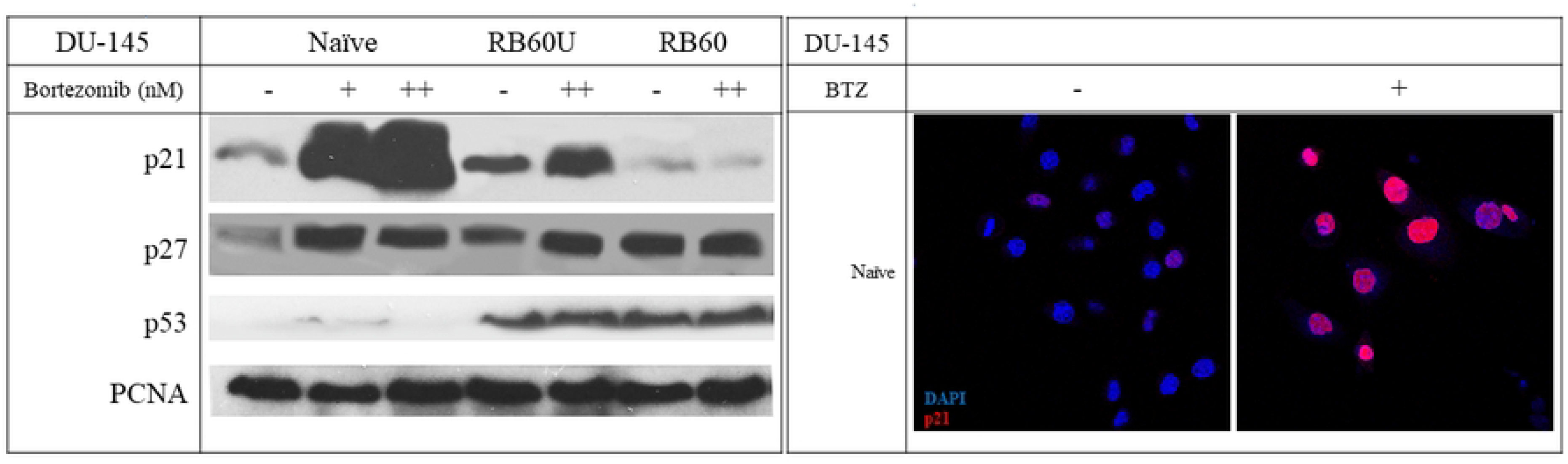
Effects of Bortezomib on main cell cycle regulators. **(A)** Western Blot analyses of key cell cycle proteins indicated accumulation of p21, p27 and p53. The experiments were conducted after 24 h of Bortezomib incubation, following a 48 h Bortezomib-deprivation of RB60 cells. The + corresponds to the low dose of 20 nM Bortezomib and the ++ to the medium dose of 60 nM. The DU-145 RB60 cells downregulate the accumulation of p21 observed on naïve cells (A1), however, the expression of p53 and p27 are high (A2 and A3). The levels of PCNA were also examined, however they did not indicate major baseline changes between naïve and resistant cells (A4). **(B)** Immunocytochemical staining of naïve DU-145 cells with antibodies against p21 and DAPI to visualize DNA content to assess the localization of p21. Following treatment with 20 nM Bortezomib, the nuclear localization of p21 was verified, indicating the pro-apoptotic role of p21 rather, than the pro-survival cytosolic presence.

A concentration of 20 nM was adequate to induce the accumulation on naïve DU-145 cells while the same was not observed on RB60 cells. The turnover rate of p21 is already known to be regulated through proteasomal degradation, a mechanism disrupted on DU-145 cells, leading to the sub sequential p21 protein accumulation. The resistant cells were able to downregulate p21 accumulation levels independently of the drug presence, and this characteristic remained after long-term deprivation of the drug, as shown on the RB60U cell clone (Fig 7A1).

The cell cycle regulator p27^Kip1^ was not found to follow the same accumulation pattern, as p21 on the resistant clone studied. After incubation with Bortezomib, the p27 protein maintained its high nuclear levels, both, on naïve and on resistant cells. Deprivation of Bortezomib for 24 h reduced the intracellular levels of p27 on DU-145 resistant cells imitating an untreated naïve cell (Fig 7A2). Since the p27 protein is degraded through the UPS, the similar protein levels detected, might indicate that the resistant clone that emerged in our study, has somehow managed to balance the pro-apoptotic signals descending from the accumulation of cell cycle inhibitors with survival-promoting molecules and antiapoptotic proteins.

The p53 protein was also assessed, to investigate potential alterations of its expression and activity. Levels of p53 protein were found elevated on the DU-145 resistant clones studied, a finding observed on both, RB60 and RB60U cells (Fig 7A3). Incubation of naïve cells with Bortezomib led to a slight increase in its presence, although the accumulation inside the resistant clones was incomparable.

Analysis of the cell cycle using PI indicated that at the basal state of naïve DU-145 cells (untreated) and DU-145 RB60 cells (treated), the cell cycle progresses undisrupted (Figs 8A, 8C and 8D). Treatment with Bortezomib causes cell cycle arrest and most of the cells exited the G_1_ phase entering G_0_, and those already on the G_2_ phase never progressed to mitosis (Fig 8B), a finding already noted by others (25,37,44,48,49). Cell cycle inhibition has already been associated with p21 and p27 accumulation, following Bortezomib treatment in vivo and the restoration of this system on the resistant clones studied surely allows survival and proliferation.

**Fig 8.**
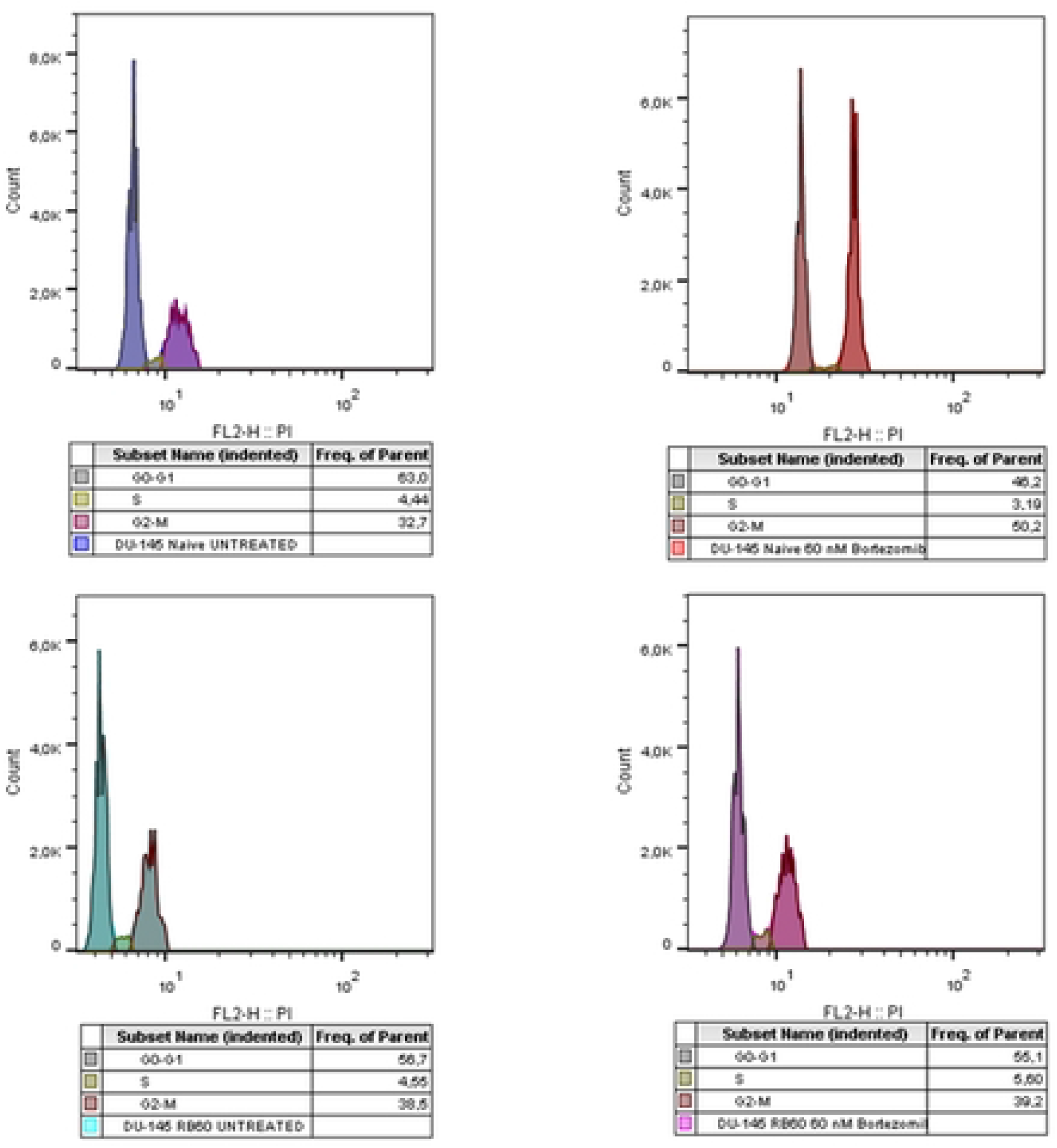
Cell cycle analysis with flow cytometry using PI. Cells were treated with Bortezomib for 24 h, following 48 h of drug deprivation from the DU-145 RB60 cells. After trypsinization, the cells were fixed using ice-cold methanol and later stained with propidium iodide. G2 arrest was observed on the treated naïve clone while the RB60 clone was found to have fully abolished Bortezomib effects. The RB60 cells did not indicate any major changes after Bortezomib treatment and had similar phase distributions with the untreated naïve clone.

Finally, the presence of PCNA was assessed using a western blot to detect whether the substitution of the cell’s ability to multiply on a normal (compared to naïve cells) rate relies on the overexpression of replication-associated molecules that may surpass, the inhibitory signaling cascades. However, PCNA showed similar expression patterns in both the experimental clones, with an expected decrease in its accumulation on DU-145 cells after 24 h of treatment with Bortezomib (*Fig 7A4*).

### JAK/STAT, Akt, and MAPK mediate the survival of the resistant clone

Following the results indicating suppression of cell cycle inhibitors and the restoration of proliferative activity on DU-145 resistant clones, we examined key signaling molecules controlling these functions upstream in the signaling cascades.

The most interesting findings concerning cell survival were those associated with MAPKs. The ERK1/2 protein kinases were found to be overexpressed on the resistant and Bortezomib-deprived clones, compared to the naïve clone (Fig 9A), and additionally, the phosphorylation patterns were greatly different. After incubation with Bortezomib, the naïve cells were found to phosphorylate ERK1/2 during the first 24 h, at low Bortezomib doses (<20 nM), however, after prolonged incubation or exposure to moderate or high doses of the drug (20-60 nM) phosphorylation was abrogated at 36-48 h. The resistant clones exhibited a different activation profile. Both MAPKs were found phosphorylated during Bortezomib incubation and their activation only declined after a remarkably high dose of Bortezomib had been added, reaching the milestone of 240 nM. Between untreated and treated DU-145 RB60 cells, no significant differences in phosphorylation were observed, indicating a permanent ERK1/2 activation, independent of Bortezomib that regulated the new equilibrium (Fig 9B).

**Fig 9.**
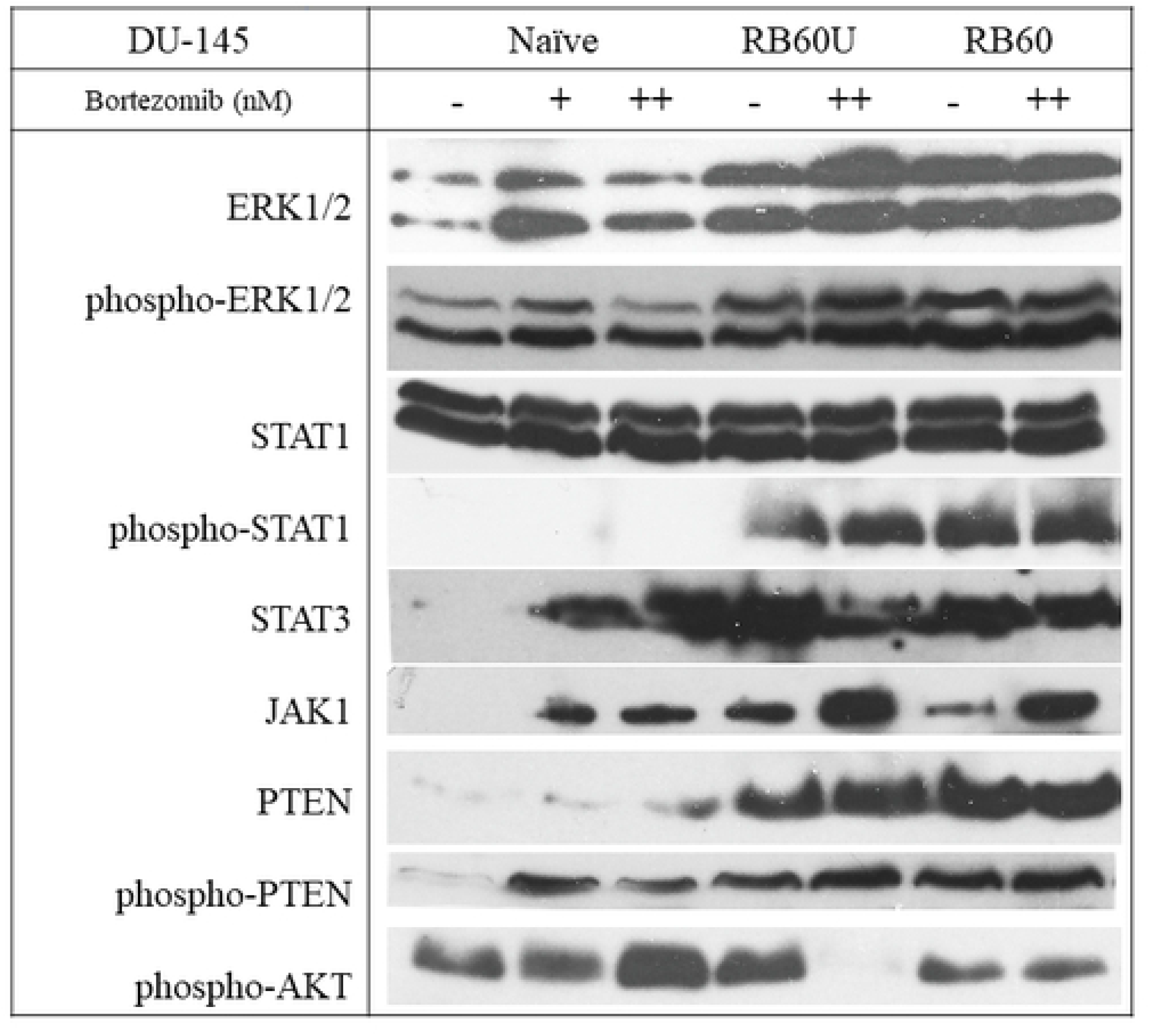
Western Blot analysis of key signaling molecules. The main pro-survival and proliferation pathways were assessed following the incubation of cells with Bortezomib. All experiments were conducted after 24 h of Bortezomib incubation following 48 h drug deprivation from the resistant clones. The cells were lysed using ice-cold lysis buffer and protein concentrations were determined using the Bradford assay. The same amounts of total protein were loaded on 12% SDS-PAGE gels and the transfer was performed using the semidry system. The + corresponds to the low dose of 20 nM Bortezomib and the ++ to the medium dose of 60 nM.

The JAK/STAT signaling pathway was also studied. JAK1 protein was found overexpressed on the DU-145 RB60 and RB60U clones, upon Bortezomib incubation (Fig 9F). JAK1 is known to phosphorylate members of the signal transducer and activator of the transcription protein family (or STAT), which facilitates the transcription of surviving-associated genes. While the total STAT1 protein was detected at similar levels among naïve and resistant cells (Fig 9C), the DU-145 RB60U and DU-145 RB60 clones had induced higher levels of phosphorylation (Fig 9D), a response pattern already connected to evasion of Bortezomib-induced apoptosis (50). The total STAT3 was also analyzed and found to be present in all cell types, however, the basal expression levels on the DU-145 naïve cells were significantly lower than those of the treated and resistant cells (Fig 9E). The JAK/STAT pathway is a major survival and differentiation pathway, active in many aggressive forms of cancer, and therefore, its activation in the resistant cells is a reasonable finding.

The activity of PTEN, a phosphatase inhibiting the AKT/mTOR pathway was also studied, as well as the activation of AKT at the key time point of 24 h. The activated forms of PTEN and AKT had opposite actions: the phosphorylated PTEN promoted apoptosis while the phosphorylated AKT favored survival. DU-145 RB60 and RB60U cells were found to express higher levels of PTEN compared to the naïve cells (Fig 9G), however, a study of its phosphorylation indicated that on the resistant clones, the dominant form is the dephosphorylated and hereby inactive (Fig 9H). The phosphorylated form of PTEN was significantly higher on the DU-145 naïve cells following Bortezomib treatment, exhibiting a dose-response pattern, while the total PTEN levels, both phosphorylated and dephosphorylated were lower at the baseline conditions.

Even though the activated PTEN is likely to induce AKT dephosphorylation, AKT was found activated following treatment with Bortezomib on the naïve cells, showing a dose-response pattern (Fig 9I). A lack of AKT activation was observed on DU-145 RB60U cells after 24 h of Bortezomib treatment, which was quite controversial, whereas the levels of phosphorylated AKT did not change on the RB60 clone. High levels of AKT have also been mentioned by other researchers (51). At the same time point, the ERK kinase is vastly phosphorylated on both, RB60 and RB60U clones, a finding indicating an alternative way to affect survival and suppression of proapoptotic signals.

### Bortezomib-resistant cells utilize autophagy as a UPS substitute

Preliminary data from other researchers have indicated an induction of autophagy in Bortezomib-resistant cells, as a balancing mechanism to obtain nutrients from damaged molecules, banish accumulated non-functional proteins and suppress the activation of pro-apoptotic pathways (25,27,31). Additionally, the joint assembly of proteasomes and autophagosomes and the emerging process called proteaphagy has been documented to increase after the administration of proteasome inhibitors, potentially as an effort to eliminate blocked proteasomes (52). To assess these phenomena, the lysosomal activity was studied using the alkaline dye Lysotracker RED, which can visualize (Fig 10C) and quantify (Fig 10B) acidic protein content. Lysotracker RED indicated increased activity inside the resistant cells, which was also quantified using flow cytometry. Additionally, by using a western blot analysis, the main autophagy biomarkers LC3 a/b, Beclin-1, and p62 were examined.

**Fig 10.**
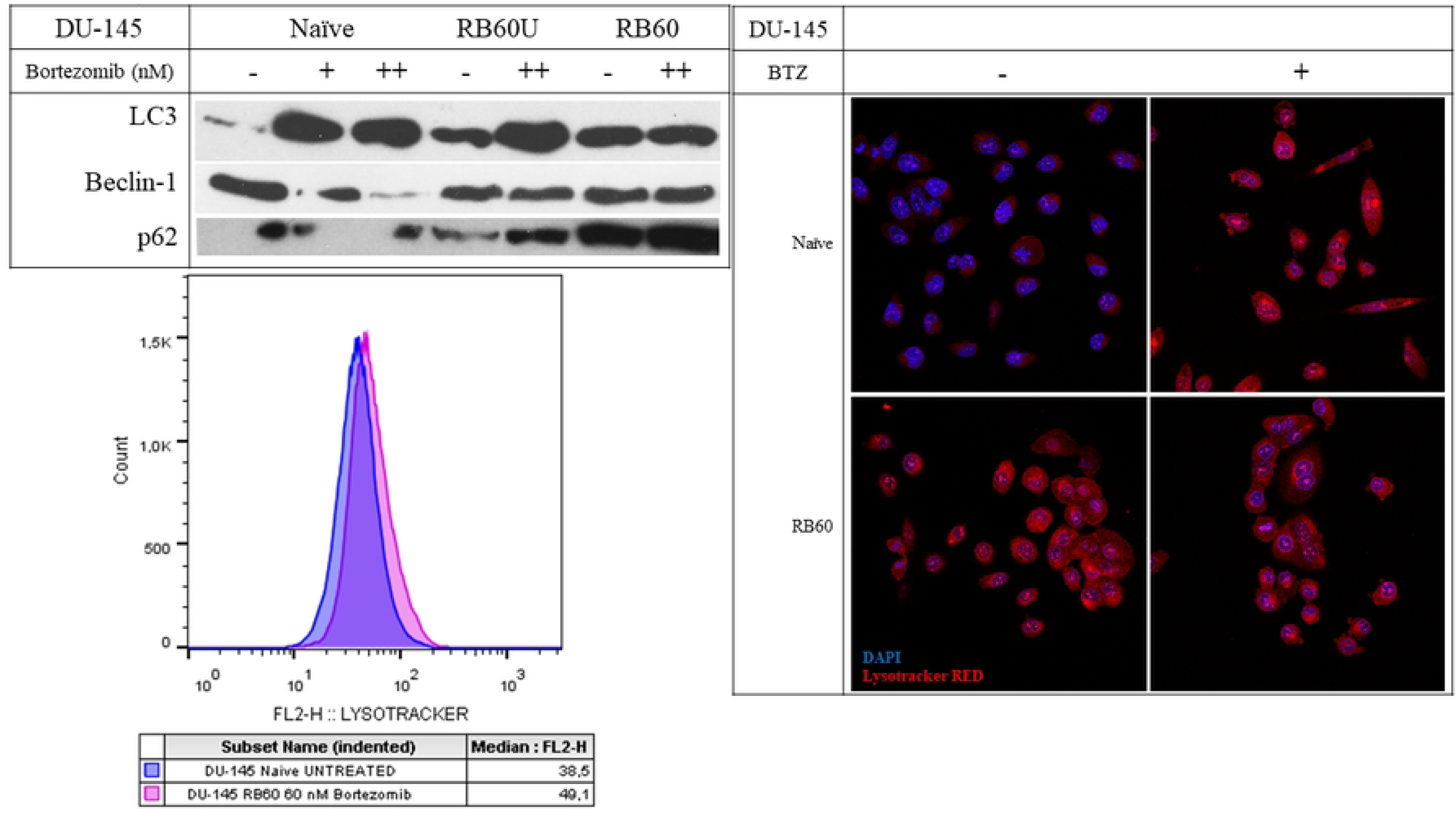
Autophagy study using Lysotracker RED and main autophagy markers using western blots. **(A)** Western Blot analyses of key proteins regulating autophagy. The experiments were conducted after 24 h of Bortezomib incubation following a 48 h Bortezomib deprivation of RB60 cells as previously noted. The + corresponds to the low dose of 20 nM Bortezomib and the ++ to the medium dose of 60 nM. **(B)** Immunocytochemical staining of naïve DU-145 cells with Lysotracker RED which stains acidic proteins, following treatment with 20 nM Bortezomib. The cells were cultured on coverslips and were stained (without fixation) with Lysotracker RED for 15 min at 37° C, followed by confocal imaging. **(C)** Flow cytometry analysis of Lysotracker RED inside naïve and resistant live cells. The cells were trypsinized and subsequently stained with Lysotracker RED for 45 min followed by analysis using a FACS Calibur flow cytometer.

The LC3 protein is a major autophagy marker. Compared to the untreated cells, the administration of Bortezomib generally increased LC3 accumulation on both, naïve and resistant clones. However, the three clones, DU-145 RB60, DU-145 RB60U, and DU-145 RB60 differed vastly at the basal levels of LC3 expression. Basal LC3 expression was the lowest in the naïve cells, and the long-term deprivation of Bortezomib on DU-145 RB60U cells led to increased LC3 levels, which were slightly lower than those of the resistant RB60 clone. Upon treatment with the drug, the naïve cells exhibited induction of LC3, while in the resistant clone, LC3 expression remained relatively stable. The RB60U clone showed an intermediate pattern between the two states, exhibiting basal LC3 levels, similar to those of the resistant clone, indicating the permanent alteration of cell functions, while at the same time exhibiting an LC3 overexpression as in the naïve clone. Autophagy seems to be regulated by the same pathways that allow the emergence of resistance since the basal autophagy levels of the RB60U and RB60 clones were almost identical (Fig 10A1).

Beclin-1 is a key molecule in the crosstalk between autophagy and apoptosis, credited with an antiapoptotic role (53,54). High expression of Beclin-1 leads to autophagy activation and its upregulation is believed to be connected to Bortezomib resistance. Knockdown of Beclin-1 has been shown to reverse Bortezomib resistance in cancer cells by inducing apoptotic cell death (34). The DU-145 naïve cells were found to have increased basal levels of Beclin-1 that declined following treatment with Bortezomib, justified by the simultaneous apoptosis induction. The DU-145 RB60 and RB60U cells maintained relatively similar levels of Beclin-1 during treatment with the drug that was higher than those of the treated naïve cells. The stable accumulation of Beclin-1 on the resistant clones upon exposure to Bortezomib explained the substitution of UPS impairment by proteophagy and the maintenance of a low apoptotic rate, similar to that of the naïve cells. A dose of 60 nM Bortezomib was found adequate to significantly downregulate Beclin-1 accumulation on naïve DU-145 cells, while the same dose on the long-deprived clone DU-145 RB60U did not provoke any changes in its presence (Fig 10A2).

p62 protein acts as a cargo receptor transporting ubiquitinated proteins to the autophagosomes and assembling structures, known as proteaphagosomes (52). During the process of autophagy, its levels decrease through subsequential degradation. The naïve cells were observed to maintain low p62 levels, even after Bortezomib treatment, showing only moderate changes. However, the resistant clones showed a different pattern. The p62 protein was overexpressed inside the resistant DU-145 RB60 and DU-145 RB60U clones, indicating long-term changes in the way cells utilize their nutrients and maintain their homeostasis (Fig 10A3). The accumulation of p62 may be the key mechanism of the autophagy upregulation, as an alternative pathway to substitute proteasome-mediated protein degradation.

### Resistant cells have lower oxidative stress levels and are less prone to drug-induced damage

Oxidative stress is a proposed mechanism of action for many chemotherapies, acting by damaging vital biomolecules and inducing cell death. Bortezomib is believed to exert a similar action, because of the disruption of cell energetics, the dysregulation of degradation pathways, and the subsequential accumulation of free radicals, resulting from the failure of the counteractive homeostatic mechanisms(9,55–58). The oxidative stress levels of naïve and resistant cells were assessed using the H_2_DCFDA assay. The resistant cells indicated lower intracellular Reactive Oxygen Species (ROS) levels compared to the naïve clone (Fig 11B) Specifically, impairment of the UPS system is believed to increase oxidative stress on cancer cells, and this effect was verified in our study, however, the resistant cells, are found to achieve lower ROS levels than those achieved by the naïve untreated cells, possibly by inducing antioxidative defense mechanisms to survive under regular doses of Bortezomib.

**Fig 11.**
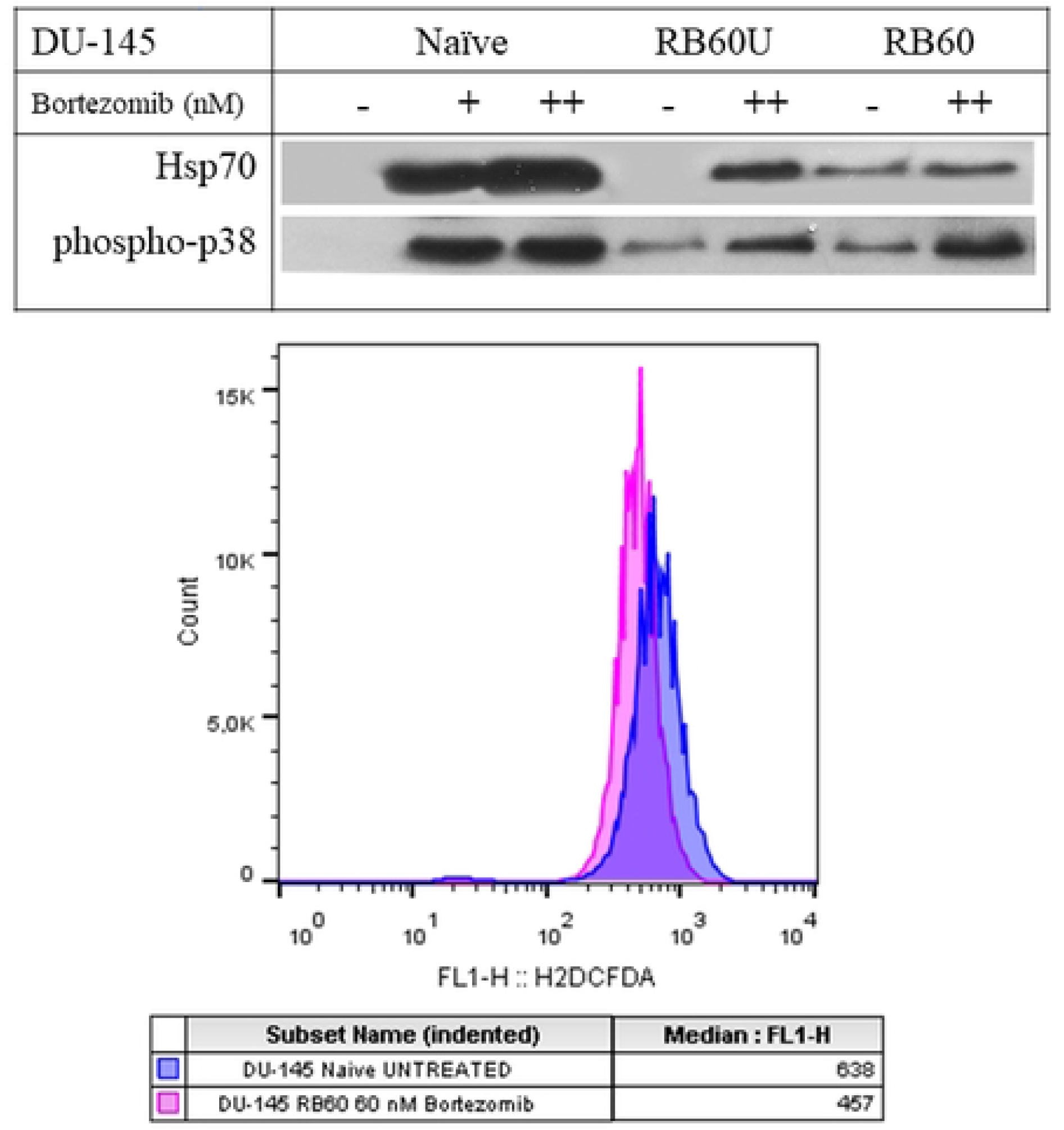
Drug-induced stress and ROS generation assay. **(A)** Western Blot analysis of phospho-p38 MAPK and Hsp70. The experiments were conducted after 24 h of Bortezomib incubation following a 48 h Bortezomib deprivation of RB60 cells. The + corresponds to the low dose of 20 nM and the ++ to the medium dose of 60 nM. **(B)** Flow cytometry analysis of ROS generation using H_2_DCFDA. The cells were incubated for 24 h with Bortezomib and then were trypsinized. During the staining procedure, they were maintained inside a culture medium to avoid heat shock and starvation stress. Staining was performed at 37° C and the samples were rinsed with PBS and analyzed using FACS Calibur flow cytometer. The resistant clone (DU-145 RB60) exhibited lower oxidative stress levels than the naïve DU-145 cells even with the presence of Bortezomib.

We further examined the effects of Bortezomib on proteins related to stress conditions, such as the heat shock proteins (59) and p38 MAPKs (60). The heat shock protein 70 (Hsp70) was analyzed, using western blots and indicated a dose-response induction in the presence of Bortezomib with a peak at 24 h. The DU-145 RB60 cells were capable to suppress its accumulation while the DU-145 RB60U cell clone exhibited a naïve-like phenotype regarding Hsp70 (Fig 11A1). The p38 MAPK belongs to a distinctive class of MAPKs that are activated through phosphorylation during stress conditions induced by radiation and drugs (61). Bortezomib induces such an effect on naïve cells, yet the resistant clone was once again immune to it, maintaining low phosphorylation levels (Fig 11A2). The two proteins mentioned, p38 and Hsp70 are thought to be correlated, with the molecular chaperon Hsp70, acting as a molecular chaperon mediating the nuclear translocation of the activated p38 (62,63).

## Discussion

The proteasome is a key functional structure in cancer cell homeostasis and the abrogation of its function by proteasome inhibitors (PIs) is a long-established therapeutic approach, already applied to several types of malignancies. However, the emergence of resistance against PIs is a major therapeutic drawback. Our study elucidates many aspects of the acquired resistance of prostate cancer cells, pointing out potential ways to target them and reverse them. Besides the approval for treatment of prostate cancer, for whom DU-145 cells are a well-established model, Bortezomib is used to treat hematological malignancies such as mantle cell lymphoma (64) and multiple myeloma (65), while at the same time, it has shown little efficacy against clonal cells of the Myelodysplastic Syndromes (MDS) (66).

The model of this study managed to acquire resistance against the PI Bortezomib and the resulting cell phenotype was suspected to have a strong genetic basis, since it remained stable for a prolonged period. The bottleneck effects caused by Bortezomib-induced massive cell deaths, combined with the dysregulation of other main signaling pathways on the surviving cells and the accumulation of somatic mutations, to the generation of a Bortezomib-resistant cell line, with increased cross-resistance to second-generation PIs such as Carfilzomib. The apoptotic rate of the newly emerged resistant cell clone gradually diminished, finally reaching those of naïve cells, and the same was observed regarding the resistance to cell cycle regulators. Three major cell cycle regulators were assessed; p21, p27, and p53, all of them already been studied previously and connected to the effects of Bortezomib treatment (4,20,67). DU-145 cells following incubation with PIs were found to dramatically increase p21 protein levels, which we observed to be downregulated on the resistant clones, while an increase of p27 and p53 was also observed, as has been previously mentioned. Although the levels of p27 remained relatively high on our resistant clones, compared to the basal state of naïve cells, in our opinion the most interesting finding of this study is related to the p53 protein, which is an important part of the p21/p53 axis (67). The p53/survivin system has already been associated with Bortezomib resistance, through the abrogation of Bortezomib-induced apoptosis in many types of cancer. Wild-type p53 leads to cells more susceptible to Bortezomib, while mutant variants of p53 have a strong association with Bortezomib resistance (68). p53 overexpression on our established resistant clone may have resulted following a mutation or the cell’s normal response to Bortezomib-induced stress and UPS impairment. The latter, mainly observed on the resistant clones may be warded off by a different mechanism, such as the assessed autophagy. Besides the high p27 and p53 expression levels on DU-145 resistant cells, their ability to enter the cell cycle was found restored. The combination of a wild-type p53 and functional p21, renders the second as a genome guardian, while in the absence of or the presence of a mutated p53, p21 can induce genomic instability (67). Our findings support this idea, pointing out that the restoration of p21 on DU-145 Bortezomib-resistant cells could reverse resistance.

The resistant cells’ ability to thrive in the presence of Bortezomib was further analyzed by looking at the key signaling molecules controlling cell survival and proliferation. Our study indicated that Bortezomib resistance in our clone heavily relies on the constant activation of MAPKs, the JAK/STAT pathway, and the activation of AKT. These molecules are already known to be active in many aggressive forms of human cancer and to counteract the proapoptotic signals induced by Bortezomib, and their exogenous inhibition could reverse the resistant phenotype (25). Indeed, the phosphorylated forms of STATs have been found to counteract the proapoptotic effects of Bortezomib in ovarian cancer (Kao, 2013) leading to the emergence of Bortezomib resistance. STAT3 has also been found to control the expression of β subunits of the 26S proteasome (69), therefore, the high STAT3 levels found in DU-145 RB60 cells could regulate the expression of *PSMB5,* to counteract the accumulation of dysfunctional proteasomal subunits, induced by Bortezomib binding. Phosphorylation of STATs, as well as of other survival-promoting transcriptional factors can be mediated through the ERK1/2, which in our resistant clones were found constantly activated. Inhibition of ERK1/2 phosphorylation by MEK inhibitors has been reported to reverse Bortezomib resistance in the SKM-1 MDS cell line (25). In parallel, activation of the AKT pathway indicated an induction on the DU-145 resistant clones. Moreover, in our study, the expression of the suppressing protein PTEN was found higher on both, DU-145 RB60 and RB60U clones, however, the dominant form of PTEN was the dephosphorylated one. High PTEN levels have already been documented as a consequence of Bortezomib treatment, eventually leading to apoptosis (70). The phosphorylated form of PTEN can stabilize p53 protein and the PTEN transcription can be induced by p53 (71), therefore, the crosstalk between p53 and PTEN is of major significance (72).

Additionally, the regulation of the other major protein degradation pathway, namely autophagy, was assessed and was found to be upregulated in the resistant cells. It has previously been shown that autophagy may be a key substitute for peptide degradation when the UPS system fails (33,34). The degradation of inhibited proteasomes at the autophagosomes is probably a key point in eliminating the blocked subunits (27) and this might be more easily induced when inhibition is irreversible with the newer PIs. In addition, the overall induction of autophagy may fulfill the need for amino acid recycling, as was shown by the constant p62 accumulation. The equilibrium between apoptosis and autophagy -which can also lead to cell death-is mediated through proteins that were found upregulated, mainly Beclin-1 and the MAPK p38, which was found to become phosphorylated, following Bortezomib treatment. p38 protein has been credited with multiple roles, among them, the regulation of autophagy (73). The increased basal levels of p38 in its phosphorylated form, combined with the induction of autophagy could imply that in our clone, p38 could function as a mediator of resistance. Thus, inhibiting p38, could -even partially-reset the cell’s ability to withstand the presence of Bortezomib.

A crucial element closely related to the many proteostasis subsystems is Hsp70, a protein that functions as a molecular chaperone. Hsp70 is believed to have a dual role in assisting both, protein refolding, as well as leading damaged, misfolded or non-functional proteins to the 26S proteasome subunit, after ubiquitin ligation (74). A permanent increase in Hsp70 accumulation could also function as a mechanism of resistance, however, such an effect was not observed in our study. Hsp70 levels exhibited great augmentation after Bortezomib treatment of naïve cells, in a dose-response manner, despite the low basal levels, observed in RB60 and even RB60U cells. Hsp70 levels of long-deprived DU-145 RB60U cells were identical to those of naïve untreated cells, while the presence of Bortezomib did not significantly alter Hsp70 expression, compared to the stable (and low) expression observed in the DU-145 RB60 clone. Induction of Hsp70 has been documented to suppress autophagy (75,76), therefore, despite its protein refolding role, given that our clone mainly achieves resistance through autophagy regulation, Hsp70 protein remains at low levels.

Besides its role in proteostasis, the Hsp70 protein has also been associated with stress conditions inside the cells. The downregulation of Hsp70 observed in the resistant cells, combined with the fact that these cells thrive in a Bortezomib-rich environment, adverted us to study oxidative stress levels. Proteasome inhibitors induced the elevation of ROS levels, resulting from damaged protein accumulation, increased need for energy to biosynthesize new proteins, replenish the damaged ones, and dysregulation of cellular homeostasis (40,58). Surprisingly, ROS levels in the resistant cells were found to be lower, even than those of the untreated naïve cells, suggesting that the resistant cells had rather fully reversed the oxidative stress damage induced by Bortezomib. These observations, combined with the finding that the resistant clones, cultured in our laboratory exhibited increased ChT-L activity, lead us to the conclusion that increased proteasome proteolytic activity can lower ROS levels, which may function as a survival mechanism for the drug-resistant cells. Stress-induced cell death is a long-suspected mechanism of action of the PIs, also explaining the increased rate of apoptosis-inducing to cancer cells, compared to normal cells, following exposure to Bortezomib (56). This observation of oxidative stress can be used as a therapeutic agent, acting synergistically with classical anticancer drugs. The administration of classic chemotherapeutic agents, such as anthracyclines and cisplatin in patients that eventually developed resistance to PIs could be resumed if inducers of cellular oxidative stress could be concurrently used as therapeutic agents. The potential depletion of the redox homeostasis system capacity would render the cells more susceptible to proteostasis impairment (77), and thus, the emergence of resistance would become less likely.

## Supporting information

***S1 Fig. Ubiquitination impairment of DU-145 RB60 cells.***

*The experiments were conducted after 24 h of Bortezomib incubation, following a 48 h Bortezomib-deprivation of RB60 cells. The DU-145 RB60 cells incubated with the high dose of 180 nM Bortezomib accumulate ubiquitinated proteins with the same rate as naïve cells treated with a low dose (20 nM)*

***S2 Fig. Apoptosis assay of DU-145 RB60U cells.***

*The DU-145 RB60U cell clone maintains the apoptosis evasion observed on the DU-145 RB60 cells after a 24-week drug deprivation with a slight increase of its apoptotic rate*.

***S3 Fig. Cell cycle analysis of DU-145 RB60U cells.***

*Following the same procedures as in naïve and DU-145 RB60 cells, the cell cycle of DU-145 RB60U cells was analyzed using PI and indicated a mild G2 arrest following incubation with 60 nM of Bortezomib for the first time after 24 weeks. The subsequent cell cycle inhibition does not result in apoptosis as has been shown by the rest experiments assessing apoptosis compared to the naïve DU-145 that within 48-72 h undergo apoptosis*.

